# High-resolution Spatiotemporal Changes in Dominant Frequency and Structural Organization During Persistent Atrial Fibrillation

**DOI:** 10.1101/2022.07.10.499456

**Authors:** Terrence Pong, Joy Aparicio-Valenzuela, Oluwatomisin Olurotimi Obafemi, Kevin J. Cyr, Cody Carlton, Calvin Taylor, Anson M. Lee

## Abstract

**Objective:** Analyze changes in frequency activity and structural organization that occur over time with persistent atrial fibrillation (AF)

**Background:** Little is known about the frequency characteristics of the epicardium during transition from paroxysmal to persistent AF. Accurate identification of areas of high dominant frequency (DF) is often hampered by limited spatial resolution. Improvements in electrode arrays provide high spatiotemporal resolution, allowing for characterization of the changes that occur during this transition.

**Methods:** AF was induced in adult Yorkshire swine by atrial tachypacing. DF mapping was performed using personalized mapping arrays. Histological analysis and late gadolinium enhanced magnetic resonance imaging were performed to determine structural differences in fibrosis.

**Results:** The left atrial epicardium was associated with a significant increase in DF in persistent AF (6.5 ± 0.2 vs. 7.4 ± 0.5 Hz, P = 0.03). The organization index (OI) significantly decreased during persistent AF in both the left atria (0.3 ± 0.03 vs. 0.2 ± 0.03, P = 0.01) and right atria (0.33 ± 0.04 vs. 0.23 ± 0.02, P = 0.02). MRI analysis demonstrated increased ECV values in persistent AF (0.19 vs 0.34, paroxysmal vs persistent, P = 0.05). Tissue sections from the atria showed increase in fibrosis in pigs with persistent AF compared to paroxysmal AF. Staining demonstrated decreased myocardial fiber alignment and loss of anisotropy in persistent AF tissue.

**Conclusions:** Changes in tissue organization and fibrosis are observed in the porcine model of persistent AF. Alterations in frequency activity and organization index can be captured with high resolution using flexible electrode arrays.

## Introduction

The pulmonary veins play an important role in atrial fibrillation (AF). Current guidelines classify pulmonary vein isolation (PVI) as first line surgical therapy for paroxysmal AF (Class IIa recommendation).(1) Unfortunately, the clinical efficacy of PVI is significantly lower in persistent AF suggesting that alternative substrates may exist which perpetuate AF outside of the pulmonary veins.(2) Prospective studies have confirmed the presence of bi-atrial AF sources emphasizing the need for ablation of patient-specific sources, in addition to PVI, to ensure freedom from AF.(3)

In the clinical setting, endocardial catheter-based contact mapping and non-invasive body surface mapping of dominant frequency (DF) have enabled the identification of alternative high-frequency sites that serve as sources of arrhythmogenic electrical activation.(4-7) However non-invasive surface mapping predominately reflects electrical activation of the overall atrial tissue. Furthermore current methods utilized in intracardiac endocardial mapping are often limited by the number of simultaneously positioned electrodes. Thus, the accurate identification of areas of high DF is often hampered by limited spatial resolution. New approaches that can effectively identify critical sites for treatment are needed.

We hypothesize that recent advances in flexible electrode arrays will allow for effective mapping of the epicardium and provide high spatiotemporal resolution characterization of differences in frequency activity that occur during AF.(8) Additionally, atrial fibrosis caused by tissue remodeling is known to result in non-uniform anisotropic impulse propagation which may perpetuate AF.(9,10) We further hypothesize that high-resolution spectral analysis may reveal changes in organization index during the transition between paroxysmal and persistent AF.

## Methods

All experiments were conducted in accordance with the guidelines of the Institutional Animal Care and Use Committee (IACUC) of Stanford University. Protocols for induction of paroxysmal AF (n = 5) and persistent AF (n = 5) in adult Yorkshire swine were used as previously described.(8,11). Briefly, paroxysmal AF was induced using rapid atrial pacing (RAP, 200 beats per minute). RAP was considered to be successful in the paroxysmal AF model if AF was sustained for more than 60 seconds, confirmed by continuous rhythm tracing. Persistent AF swine underwent cardiac pacemaker insertion (Assurity MRI Pacemaker, Abbott, Chicago, IL) with leads implanted into the right interatrial septum and right atrial free wall, and animals were rapidly paced at 350 bpm to induce AF, Fig. 1a. After self- sustained persistent AF was confirmed at 6 weeks, the pacemakers were turned off and the animals were maintained in persistent AF for an additional 4 weeks before terminal electroanatomic mapping. Daily oral aspirin and digoxin was administered for stroke prevention and to prevent development of clinical heart failure during high-rate atrial pacing until the terminal procedure.

**Figure 1.**
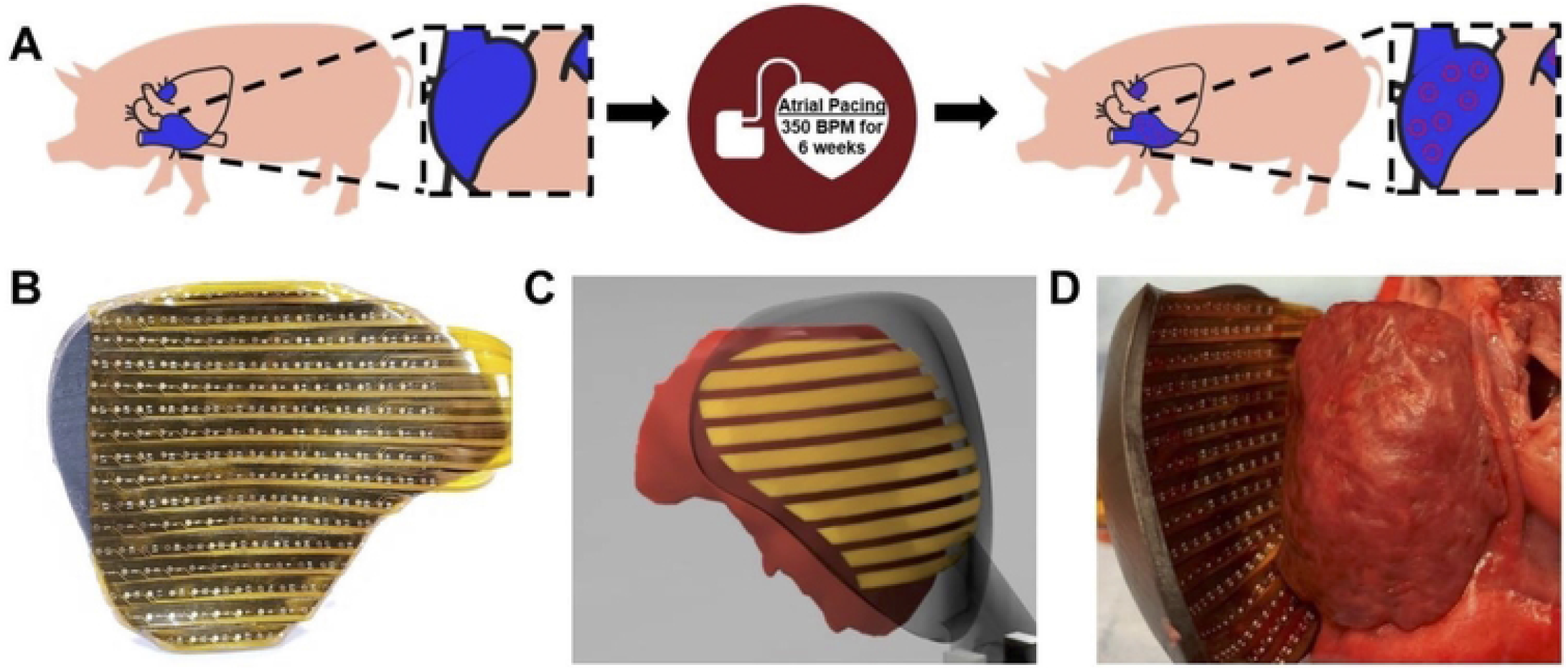
Persistent AF experimental design and electroanatomic mapping. (A) Persistent AF swine underwent cardiac pacemaker insertion and animals were rapidly paced at 350 bpm for six weeks to induce AF. (B) Flexible epicardial mapping devices were custom fabricated from animal specific cardiac magnetic resonance imaging. (C-D) Custom gold-plated flexible electrode arrays were mounted onto 3D printed epicardial shells to create a functional mapping device.

### Epicardial Mapping studies

Flexible epicardial mapping devices were custom fabricated from animal specific segmentations of the right and left atrium derived from magnetic resonance imaging. Contoured shells matching the atrial epicardial surfaces were then generated in computer aided design (CAD) software (Fusion 360, AutoDesk, San Rafael, CA) and 3D printed with a stereolithographic printer using flexible photopolymer resin (Form3, Formlabs, Somerville, MA). After printing, the devices were UV-cured and custom gold-plated flexible electrode arrays were mounted onto the epicardial shell to create a functional mapping device, Fig. 1b-d. The final epicardial mapping device consisted of up to 256 unipolar electrodes with 3 to 4 mm interelectrode spacing optimized to cover the global epicardial surface.

Prior to terminal mapping, each animal was premedicated with Telazol, Ketamine, and Xylazine; intubated; and anesthetized with isoflurane. Surface ECG and arterial blood pressure recordings were continuously monitored. A median sternotomy was performed, and the epicardial mapping devices were placed directly onto the heart to collect electroanatomic mapping data. Multiple 60 second epochs of AF were recorded per animal. All electrogram data was recorded using a biopotential measurement system (2048Hz sample rate, BioSemi, Amsterdam, Netherlands) and exported to MATLAB (Mathworks, Natick, MA) for data analysis.

### Histological Staining

Tissue samples obtained from the endocardium and epicardium were embedded in OCT cryo-embedding media and sliced in 15um thick slices. The sections were stained with H&E and Masson’s Trichrome. Actin filaments were stained with Phalloidin. Images were obtained with a field microscope at 10x and a LSM880 inverted confocal microscope at 40X. Analysis of fibrosis percentage was performed using Computer assisted histomorphometric analysis software in Python. The software estimates cardiac muscle (red stained areas) and connective tissue (blue stained areas) and computes the ratio of muscle and connective tissue.

### MRI analysis

Prior to mapping, the pigs were sedated, intubated, and taken to the MRI scanner. The scanner used was a Signa HDx 3.0T (GE Healthcare). We used an 8-channel chest coil. The sequences we used for the pig in vivo studies were, FIESTA (SAX, LAX cine), FGRE-IR (LGE), SMART1, MOLLI (T1 mapping), 3D-MDE (3D LGE, early/late phase). For contrast, we injected 0.2mmol/kg of Multihance, a Gadolinium based contrast agent (Gd).

Obtained SMART1 and MOLLI sequences were processed using the qMRLab Software package for MATLAB (12). The Barral Method for Inversion Recovery was used to generate T1 maps from image sequences (13). From generated T1 maps, pixel-by-pixel T1 values were compiled into a histogram, and the most common T1 value selected.

The medial atrial wall was isolated using hand-drawn image masks for T1 map generation. The atrial wall was manually traced and reconstructed with axial cuts for the analysis. This region was selected for its visibility and stability on MRI, in addition to previous evidence of AF-related fibrosis (14). T1 maps of blood were generated from the contents of the right ventricle for convenience. Myocardial fibrosis was quantified using extracellular volume fraction (ECV), calculated from pre and post contrast T1 relaxation times using the Gd partition coefficient (15). All pigs were healthy aside from induced AF; therefore hematocrit was estimated for all pigs to be 40%.

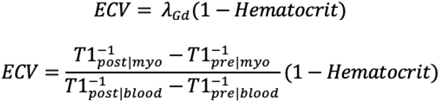

## Data Analysis

Electrogram data were processed in a custom-designed MATLAB program. Unipolar electrograms were bandpass filtered (1 – 400Hz) and far-field ventricular QRS-T signaling was subtracted before atrial signal analysis. Activation times were determined by (dV/dt) _max_ and manually reviewed and edited for accuracy. Channels with poor contact or noise were removed, and data interpolated. Isochrone activation maps depict activation times of the entire mapped atrial epicardium, with red indicating early activation and blue indicating late activation. Power spectrum analysis was performed with additional bandpass filtering (between 3 and 40Hz), rectification, and Hanning windowing. A fast Fourier transform was performed over seven consecutive sliding eight-second window, with four second overlaps between windows. The frequency band with the highest strength was called the dominant frequency (DF), and DF outside of the regularity indices (RI), 0.2 were discarded. (5,16) Organization index (OI) was calculated by dividing the area under the DF and its harmonics by the total power of the frequency spectrum (17). Final maps of activation time, DF, and OI were interpolated onto 2D reconstructions of the epicardial mapping probes to aid in visual reconstruction of electroanatomic maps.

## Results

Of the paroxysmal swine group, one animal was unable to be tachypaced into sustained atrial fibrillation and was excluded from the study. Similarly, one persistent animal was excluded because of pacemaker/lead problems that failed to produce self-sustained AF at six weeks. The remaining persistent AF pigs were in AF at the time of their mapping procedure. Paroxysmal pigs were mapped after successful AF initiation with burst atrial pacing. No persistent AF animals developed signs of clinical heart failure at the time of terminal mapping.

Each flexible mapping array was designed to optimize contact with the epicardial surface of the right and left atria. However, swine possess smaller posterior left atrial walls as a result of pulmonary veins that drain into the left atrium through two ostia in a posterocranial fashion.(18) Thus, anatomical differences between humans and swine prohibited safe in vivo mapping of the posterior pulmonary veins. In light of these anatomical differences, electrograms were obtained from the atrial free walls.

### Left Atrial Dominant Frequency and Organization Index

Representative atrial electrograms obtained from mapping the left atrium of paroxysmal and persistent AF animals after ventricular-subtraction are shown in Figure 2A, B. Paroxysmal AF animals had comparatively more organized baseline activity compared to persistent AF. Figure 2C, D show example frequency spectra derived from the respective electrograms. Paroxysmal AF was characterized by more organized activity as evident by the presence of strong harmonics in Fig. 2C. Conversely, harmonic frequencies were less distinct in persistent AF electrograms, Fig. 2D. Overall, there was a significant increase in mean DF from paroxysmal to persistent AF (6.5 ± 0.2 vs. 7.4 ± 0.5 Hz, P = 0.03), Fig. 2E.

**Figure 2.**
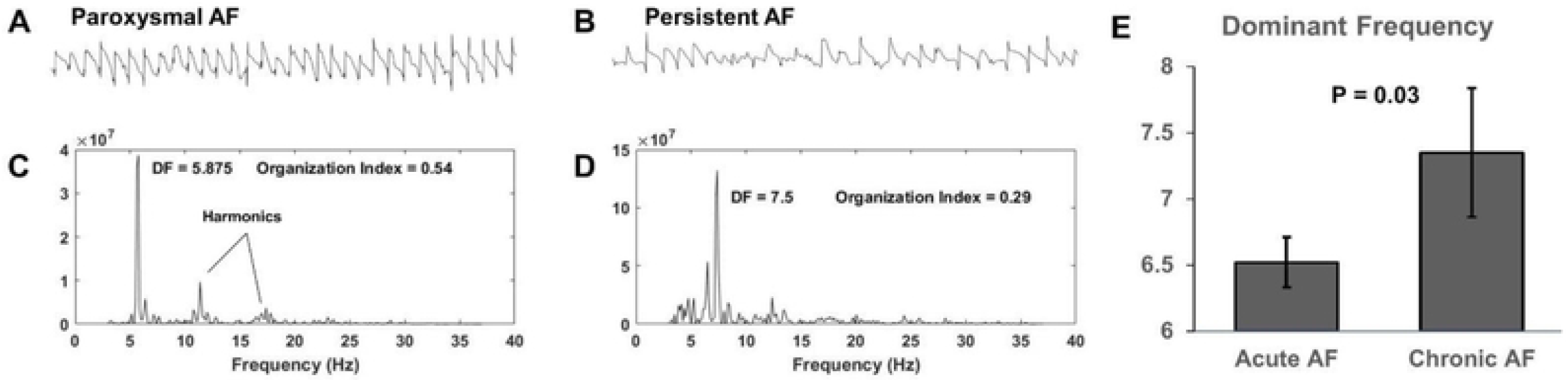
Persistent AF increased left atrial dominant frequency. Left atrium electrograms of (A) paroxysmal and (B) persistent AF. Paroxysmal AF animals had comparatively more organized baseline activity compared to Persistent AF. (C-D) Example frequency spectra derived from the respective electrograms, paroxysmal AF was characterized by more organized activity as evident by the presence of strong harmonics. Conversely, harmonic frequencies were less distinct in persistent AF electrograms. (E) There was a significant increase in mean DF from paroxysmal to persistent AF.

Figure 3 shows examples of static dominant frequency maps of the left atrium. Two-dimensional spatial mapping of DF identified areas of higher DF (in red) and lower DF (in blue). The paroxysmal AF left atrium had a larger distribution of low-DF (blue) areas as depicted in Fig. 3A, versus persistent AF in Fig. 3B. Representative electrograms for paroxysmal and persistent AF derived from the dashed boxes are shown in Fig. 3C, D. Corresponding isochrone maps of action potential wave propagation captured multiple sites of action potentials originating from sites of high DF, Fig. 3E, F.

**Figure 3.**
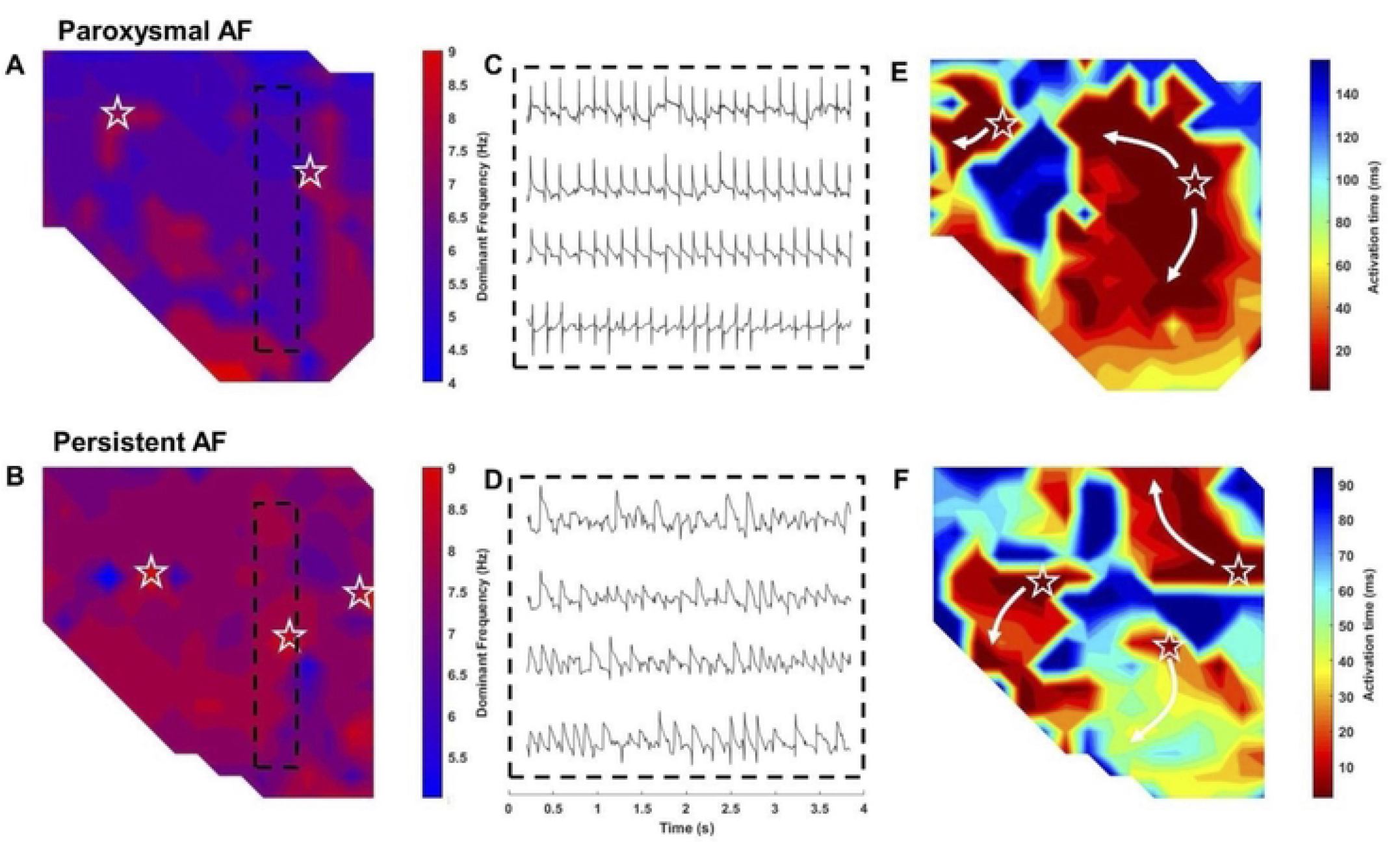
High resolution spatial mapping of dominant frequency. Two-dimensional spatial mapping of DF identified areas of higher DF (in red) and lower DF (in blue). The (A) paroxysmal AF left atrium had a larger distribution of low-DF (blue) areas, versus (B) persistent AF. (C-D) Representative electrograms for paroxysmal and persistent AF derived from the dashed boxes. (E-F) Corresponding isochrone maps of action potential wave propagation captured multiple sites of action potentials originating from sites of high DF. (☆ – denotes sites of activation).

Next, we analyzed the variability in left atrial frequency between paroxysmal and persistent AF by characterizing the organization index for individual electrograms. Figure 4A, B show example two-dimensional spatial OI maps for paroxysmal and persistent AF, respectively. There were notable clusters of similar OI that were present in paroxysmal AF that were not present in persistent maps. Persistent AF was associated with decreased left atrial OI compared to paroxysmal AF, (0.23 ± 0.03 vs. 0.30 ± 0.03, P = 0.01), reflecting the more complex AF dynamics seen in persistent AF, Fig. 4C.

**Figure 4.**
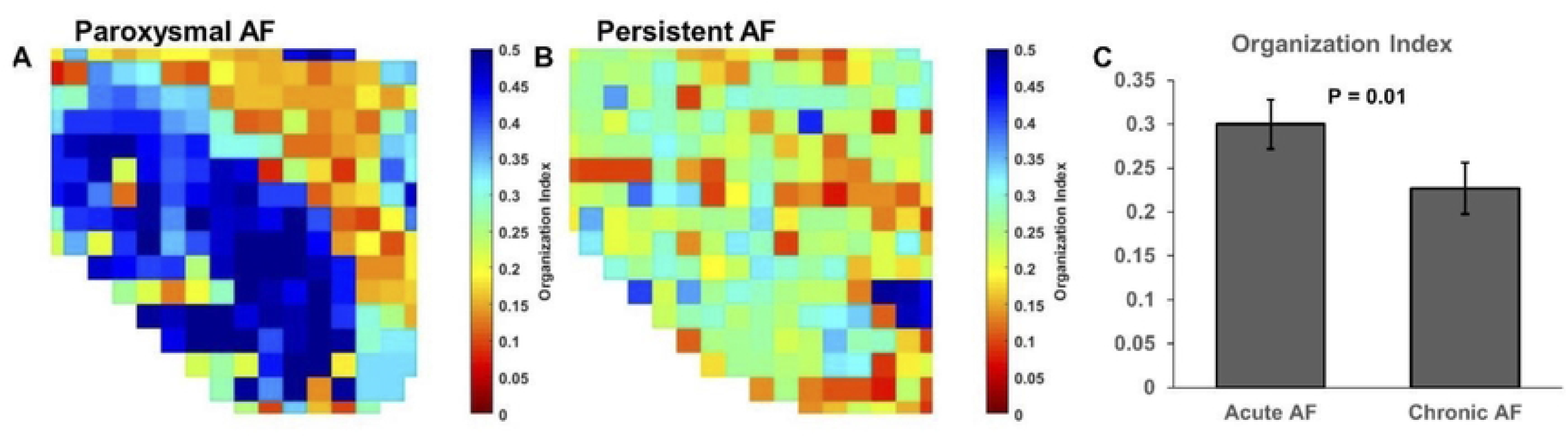
Persistent AF decreases left atrial organizational index. (A-B)Example two-dimensional spatial OI maps for paroxysmal and persistent AF, respectively. There were notable clusters of similar OI that were present in paroxysmal AF (grouped blue and orange areas) that were not present in persistent AF maps. (C) Persistent AF was associated with decreased left atrial OI compared to paroxysmal AF.

### Right Atrial Dominant Frequency and Organization Index

We performed similar DF and OI analysis for paroxysmal and persistent AF electrograms obtained from the right atrium. Figure 5A,B show representative DF maps for paroxysmal and persistent AF respectively. There was no significant difference observed in right atrial DF in respects to with persistent AF (7.0 ± 0.5Hz), and paroxysmal AF (6.9 ± 0.6Hz, P = 0.72), Fig. 5C. However, there was a significant change in OI with persistent AF, as seen in representative OI maps, Fig. 5D, E. The OI significantly decreased during persistent AF in the right atria (0.23 ± 0.02 vs. 0.32 ± 0.04, P = 0.02). In general, we observed a greater degree of anatomic variability in right atrial morphology compared to the left. While persistent AF tended to have a higher DF in the left atrium than the right, our study did not show significance.

**Figure 5.**
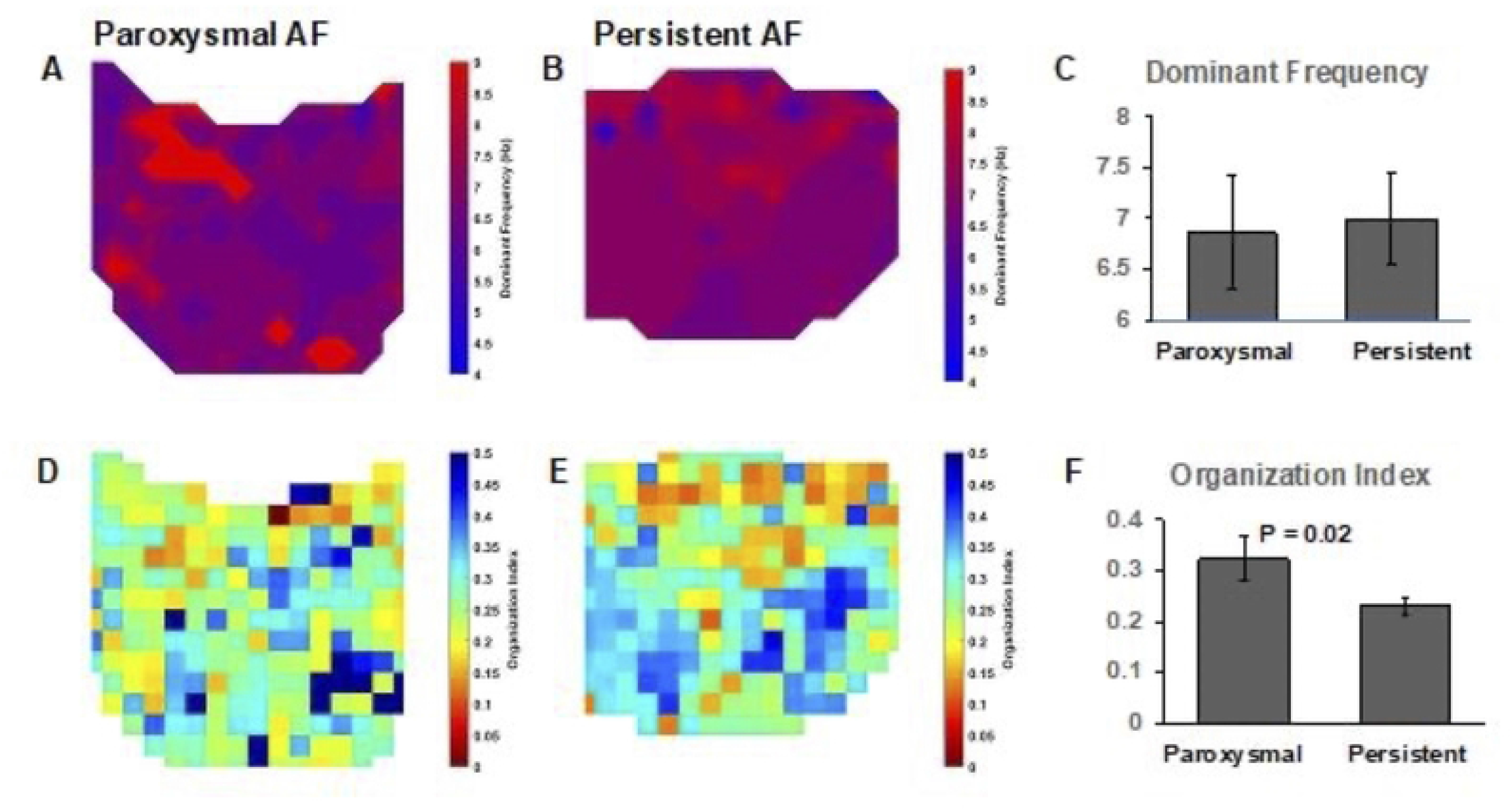
Right atrial changes in dominant frequency and organizational index. (A-B) Representative right atrial DF maps for paroxysmal and persistent AF. There was no significant difference with the right atrial DF between paroxysmal andpersistent AF (6.9 vs. 7.0, P = 0.72). (C,D) Representative right atrial OI maps of paroxysmal and persistent AF. Right atiral OI significantly decreased during persistent AF (0.23 vs. 0.32, P = 0.02).

### Histomorphometrical and MRI Analysis of Endocardial and Epicardial Fibrosis

Endocardial and epicardial tissue sections taken from the left and right atrium were stained with Hematoxylin and Eosin, Masson’s-Trichrome, and Phalloidin to compare the degree of fibrosis between paroxysmal and persistent AF. As shown in Fig. 6, both the endocardium and epicardium experienced an increase in fibrosis after six weeks of persistent AF, Fig.7. Analysis of Masson’s-Trichrome stains revealed a significant increase in epicardial fibrosis in persistent AF (3.1% vs 11.6%, paroxysmal vs. persistent AF, P = 0.01). Staining of the endocardium demonstrated a similar result; increased fibrosis in persistent AF (1.9% vs. 3.2%, P = 0.05). Phalloidin staining for F-actin demonstrated decreased myocardial fiber alignment and loss of anisotropy in persistent AF tissue.

**Figure 6.**
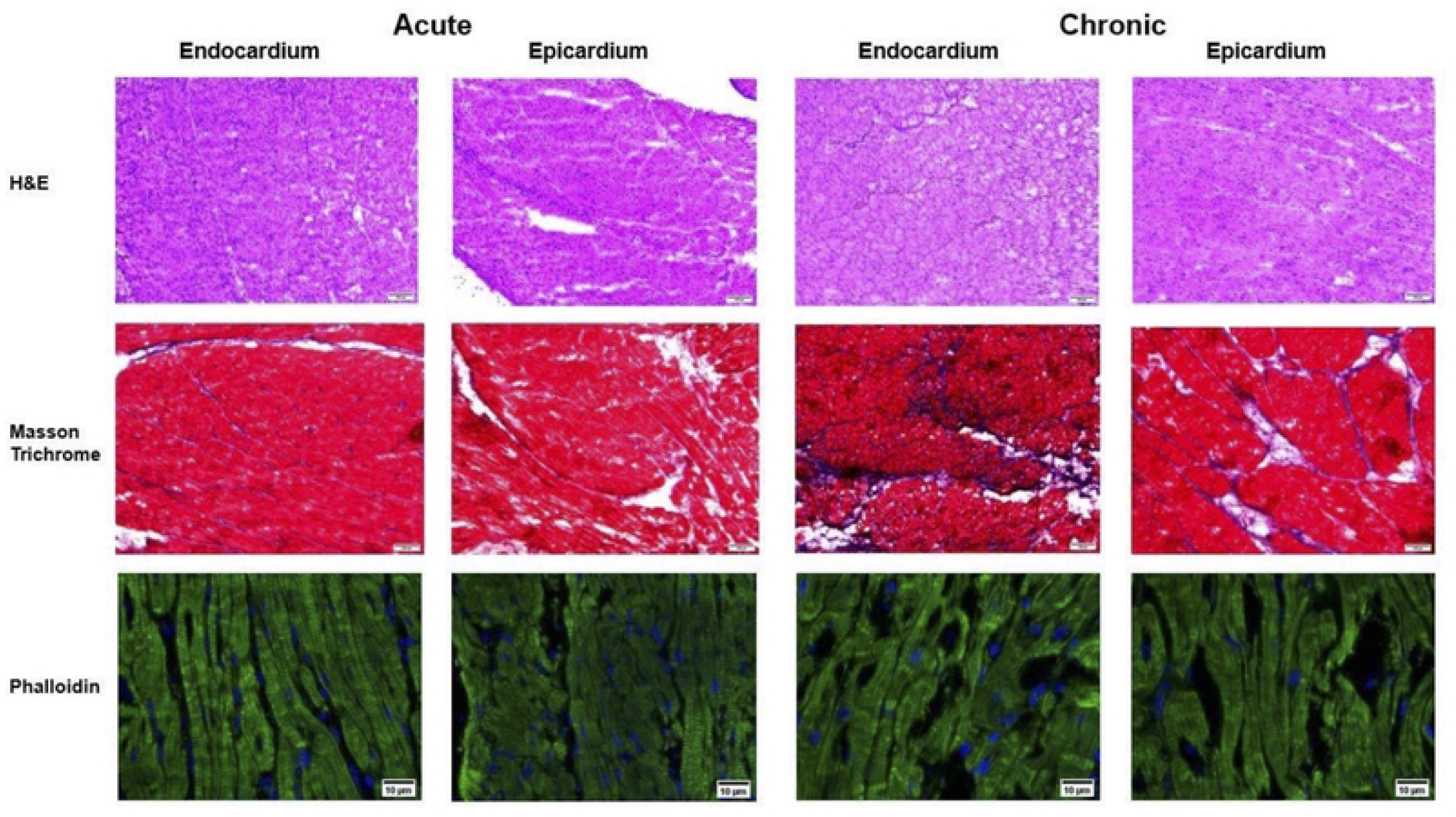
Histological analysis of atrial fibrosis and actin filament alignment. Endocardial and epicardial tissue sections taken from the left and right atrium were stained with Hematoxylin and Eosin, Masson’s-Trichrome, and Phalloidin to compare the degree of fibrosis between paroxysmal and persistent AF. Both the endocardium and epicardium experienced an increase in fibrosis after six weeks of persistent AF. Phalloidin staining for F-actin demonstrated decreased myocardial fiber alignment and loss of anisotropy in persistent AF tissue. Scale bar 50um.

**Figure 7.**
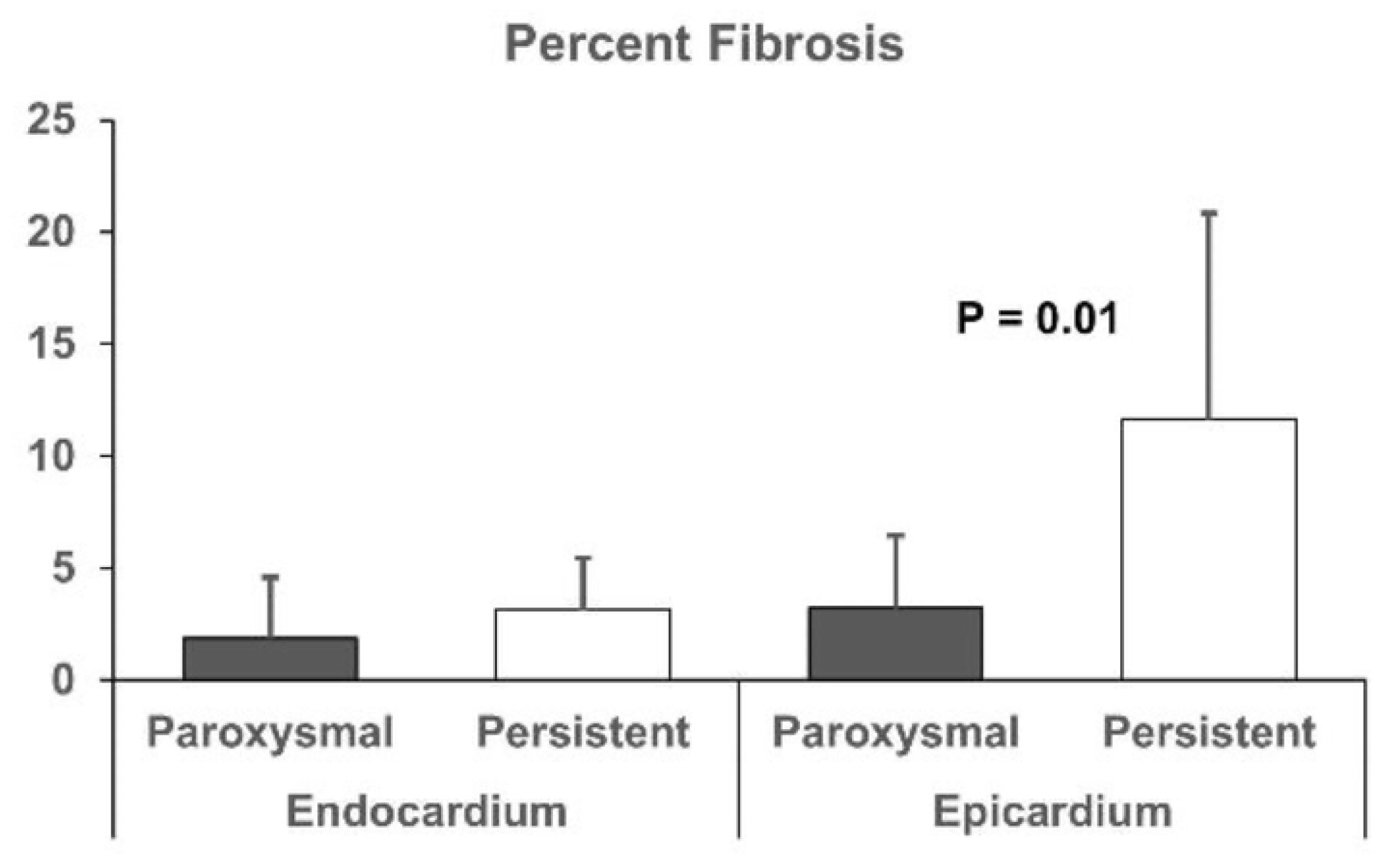
Persistent AF causes increased epicardial fibrosis. Analysis of Masson’s-Trichrome stains revealed a significant increase in epicardial fibrosis

CMR myocardial Fibrosis quantification calculations yielded expected results. Pre and post Gd T1 relaxation times of paroxysmal and persistent AF pigs were used to calculate ECV. The average pre-Gd T1 relaxation time of myocardium for paroxysmal Af pigs was 1,133.3333 ±87.451 and post-Gd T1 relaxation time of 561.6667 ±53.94

In comparison, the average T1 relaxation time of myocardium for persistent AF pigs pre-Gd was 1,315 ±101.949 and 386.6667 ±93.009 or post-Gd T1 values. Variation in T1 values were within expectations from previous research. Results showed a trend of higher ECV values for persistent AF pigs in comparison to paroxysmal AF (0.19 vs 0.34, paroxysmal vs persistent, P = 0.05), implying a greater extent of tissue fibrosis, Fig. 8.

**Figure 8.**
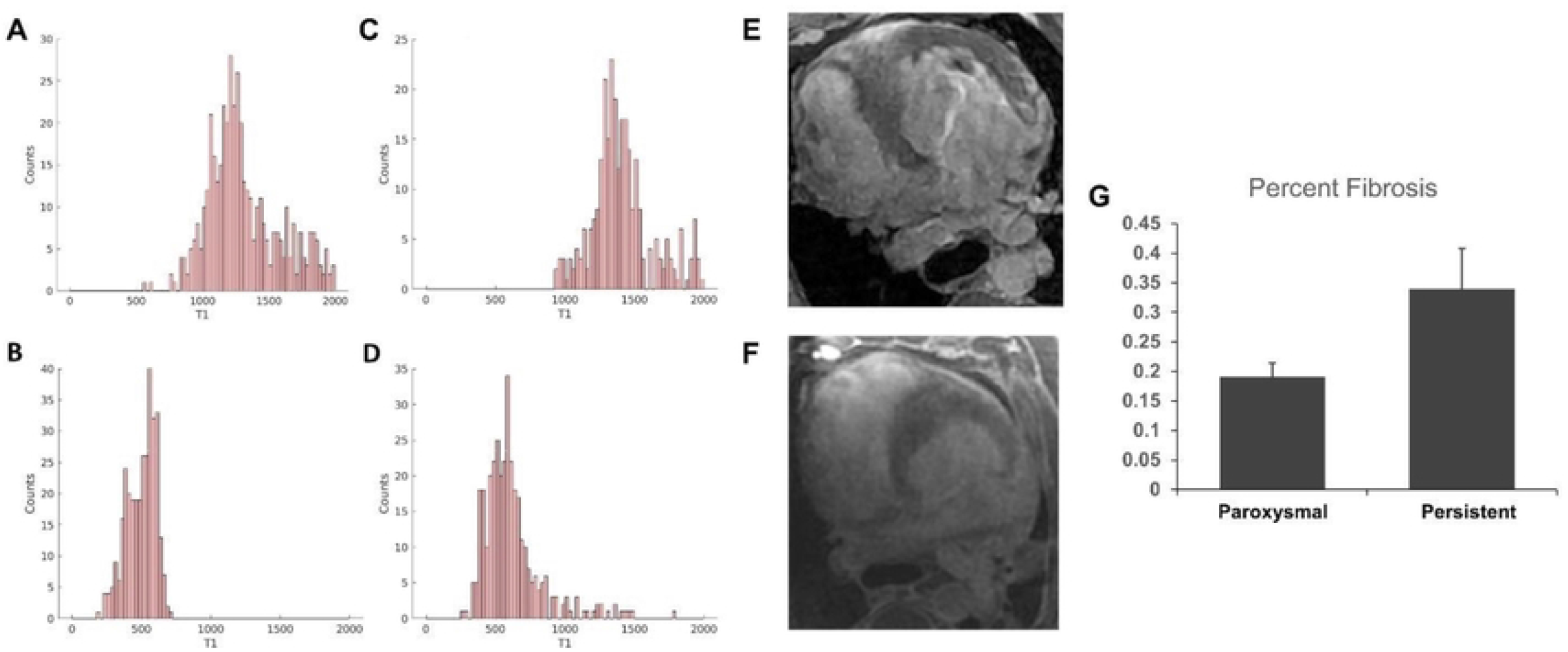
Cardiac fibrosis quantification by magnetic resonance imaging in pigs. Example T1 relaxation times of paroxysmal AF (A) pre-gadolinium and (B) post-gadolinium MRI imaging. Similar analysis of T1 relaxation time in Persistent AF of (A) pre-gadolinium and (B) post-gadolinium showed an average increase of 181.67ms (pre-Gd), and average decrease of 175ms (post-Gd). (E) Persistent AF MRI demonstrating left atrial fibrosis compared to (F) Paroxysmal AF MRI demonstrating absence of left atrial fibrosis. (G) ECV calculation demonstrated an increased level of tissue fibrosis in persistent AF compared to paroxysmal AF (p = 0.05).

## Discussion

Using a porcine model, we have demonstrated myocardial structural remodeling and increased tissue fibrosis with persistent AF. Novel flexible electrode arrays were employed to collect electrogram data from the epicardial surface and capture, in high-resolution, changes in atrial organization index and dominant frequency. Persistent AF resulted in a decrease in OI and an increase in the DF. Thus, physical changes in the form of decreased tissue organization and increased myocardial tissue fibrosis may contribute to changes in atrial electrophysiological activity during persistent AF.

Studies of myocardial fibrosis during atrial fibrillation suggest that pathological tissue remodeling results in progressive electrophysiological and structural changes that promote and perpetuate atrial fibrillation (19-22). Changes in cytoskeletal architecture have been shown to directly affect electromechanical cardiomyocyte dynamics (23). Ultrastructural changes that occurred in the setting of persistent AF were characterized by histomorphometrical analysis and MRI analysis. We observed decreased myocardial fiber alignment and loss of anisotropy in actin filament alignment with persistent AF. On a macro level, characterization of atrial fibrosis through MRI ECV estimation showed that persistent AF pigs had a greater extent of tissue fibrosis.

The relationship between ultrastructural changes and electromechanical atrial remodeling has long been postulated to affect clinical function and recovery after direct cardioversion (24,25). Persistent AF leads to pathologic remodeling that has a direct impact on the atrial conduction system. Studies have shown that atrial defibrillation during periods of high AF organization may increase shock efficacy, and that atrial defibrillation shocks delivered during periods with high OIs (> 0.5) correlate with improved defibrillation efficacy (26). Our high-resolution maps show that persistent AF is associated with decreased OI, as well as lost of organizational clustering seen in paroxysmal AF.

The decreased efficacy of PVI in persistent AF might then be explained by the development of high DF foci outside of the region of pulmonary veins. These foci are likely the result of decreased OI and pathologic remodeling that affects the conduction system

It is estimated that 12.1 million people in the United States will have atrial fibrillation in 2030 (27). There remains much to learn about the disease process that may yet lead to advances in therapy and management. Technological advances improve our ability to collect data. Recent advances in spatiotemporal resolution allows for better characterization of the changes in frequency activity that occur during AF. We anticipate that further advances in technology will continue to improve spatiotemporal resolution and characterization of frequency changes that occur with AF. These developments will allow for improved therapies to better treat persistent AF.

## Acknowledgements

Animal experiments were performed in Stanford shared facilities supported by NIH S10RR029020-01.

## Author contributions

TP and AML designed the research study; TP and OOO wrote the paper; TP, OOO, JAV, KJC, and CC conducted the experiments and statistical evaluations; OOO and CT performed MRI fibrosis analysis. All authors were responsible for final edit and approval of the manuscript.

